# Unraveling the Interplay between Phytochemicals and Rhizosphere Bacteria in *Ficus racemosa*

**DOI:** 10.1101/2024.02.27.582418

**Authors:** Acharya Balkrishna, Priya Kaushik, Sourav Ghosh, Vedpriya Arya

## Abstract

Plant-microbe interactions in the rhizosphere play crucial roles in plant health and ecosystem functioning. Despite extensive research, gaps persist in understanding the relationship between phytochemicals and rhizosphere bacteria, particularly in species like *Ficus racemosa*. This study aims to investigate this interplay by characterizing the diversity and abundance of rhizosphere bacteria in *Ficus racemosa* and assessing the influence of phytochemical composition on microbial community structure and function. Through a multidisciplinary approach integrating microbiology and phytochemistry, our findings contribute to a holistic understanding of rhizosphere ecology and inform strategies for optimizing plant-microbe interactions in agricultural and environmental contexts.

## Introduction

Plant-microbe interactions in the rhizosphere, the narrow region of soil surrounding and influenced by plant roots, are pivotal for shaping plant health, growth, and ecosystem functioning. In this microenvironment, a dynamic interplay occurs between plants and various microorganisms, including bacteria, fungi, and archaea, resulting in symbiotic, pathogenic, or commensal relationships that significantly impact plant nutrition, stress tolerance, and disease resistance.

Symbiotic interactions are particularly noteworthy, exemplified by mutualistic relationships between plants and beneficial microbes. For instance, leguminous plants engage in symbiosis with nitrogen-fixing bacteria like Rhizobium spp. and Bradyrhizobium spp., enhancing nitrogen availability and promoting plant growth by converting atmospheric nitrogen into a usable form. Similarly, mycorrhizal fungi form mutualistic associations with most land plants, facilitating nutrient uptake, particularly phosphorus and micronutrients, in exchange for organic carbon compounds.

Moreover, the rhizosphere harbors diverse plant growth-promoting rhizobacteria (PGPR) that bolster plant health through various mechanisms, including phytohormone production, nutrient solubilization, and disease suppression. However, some rhizosphere microbes can be detrimental, causing diseases that impair plant growth and yield. Understanding the dynamics of these interactions is crucial for devising sustainable disease management strategies in agriculture. Factors such as soil type, plant species, root exudates, and environmental conditions influence the composition and functioning of the rhizosphere microbiome. Advances in high-throughput sequencing technologies have unveiled the complexity of microbial communities in the rhizosphere and their roles in plant-microbe interactions.

Despite significant research in this area, there’s a notable gap concerning the specific relationship between phytochemicals—bioactive compounds produced by plants—and rhizosphere bacteria, especially in species like *Ficus racemosa*. Existing studies often focus on individual aspects without fully elucidating the interconnectedness between phytochemicals and microbial communities or their implications for plant health and ecosystem resilience. Therefore, this study aims to explore the relationship between phytochemicals and rhizosphere bacteria in *Ficus racemosa*. By characterizing the diversity and abundance of rhizosphere bacteria and investigating the influence of phytochemical composition on microbial community structure and function, we seek to unravel the mechanisms driving plant-microbe interactions. Through a multidisciplinary approach integrating microbiology and phytochemistry, our findings aim to contribute to a holistic understanding of rhizosphere ecology and inform strategies for optimizing plant-microbe interactions in agricultural and environmental contexts.

## Material Methods

### Site selection and Plant selection

When selecting plants for cultivation or conservation, their occurrence in a specific geographic region is a crucial consideration. This criterion ensures species adapt and thrive in the local environment. Focusing on indigenous plants leverages their inherent ability to flourish in prevailing climate, soil conditions, and ecosystem. Through meticulous assessment of medicinal plant occurrences, stakeholders can make informed decisions, preserving regional flora, promoting ecosystem health, and harnessing plants adapted to unique ecological niches. Ten multirange medicinal plants were carefully selected for study based on authoritative sources and field observations (Table1). These plants offer valuable insights into medicinal properties and adaptability to diverse ecological conditions. This approach not only ensures ecological sustainability but also preserves regional flora and maximizes the benefits of native plants. To support this study, we have referred to various authoritative sources such as the Flora of British India (1885), Flora of Upper Gangetic Plain (1960), Botany Bihar Orissa (1961), Flowering plants of Uttarakhand, a Checklist (2007), Flora of Uttar Pradesh (2020 & 2022), and conducted field observations.

**Table 1.** Medicinal *Ficus racemose* with their medicinal property.

| Name of the Selected Plant | Common and Vedic name | Type | Family | Ayurvedic | Medicinal Uses | Pharmacological activities | Reference |
| --- | --- | --- | --- | --- | --- | --- | --- |
| *Ficus racemosa* L. | Ambar, Domoor, Dumar, Gular, Phagoora, Udumbara, Umar, Umba (Hindi); पयोवृक्षक: उदुम्बरः(Payovrkṣakaḥ udumbarah) | Tree | Moraceae | yes | Diabetes, liver disorders, inflammatory conditions, hemorrhoids, respiratory and urinary diseases. | Anti-inflammatory | (Balkrishna A., (2022) |

### Occurrence of Plants in the Present Study

The selection of plants for this study involves a thorough process starting with an analysis of the state’s climate, soil conditions, and native vegetation. It’s crucial to understand the unique ecosystems within the region and identify plants adapted to local conditions. Factors such as plant hardiness zones, site-specific characteristics, and potential microclimates further guide the selection process. Collaboration with local authorities and experts, including botanical gardens or agricultural extension offices, provides valuable insights. Prioritizing native species fosters biodiversity and ecological balance, while guarding against invasive species helps preserve a healthy ecosystem. Tailoring choices to urban or garden settings, promoting diversity, and conducting small-scale trials contribute to successful plant selection. Documentation and ongoing observation aid in learning and improving future landscaping efforts for a sustainable and thriving plant environment. Table 2 lists the occurrence of Ficus racemose based on their distribution.

**Table 2.**
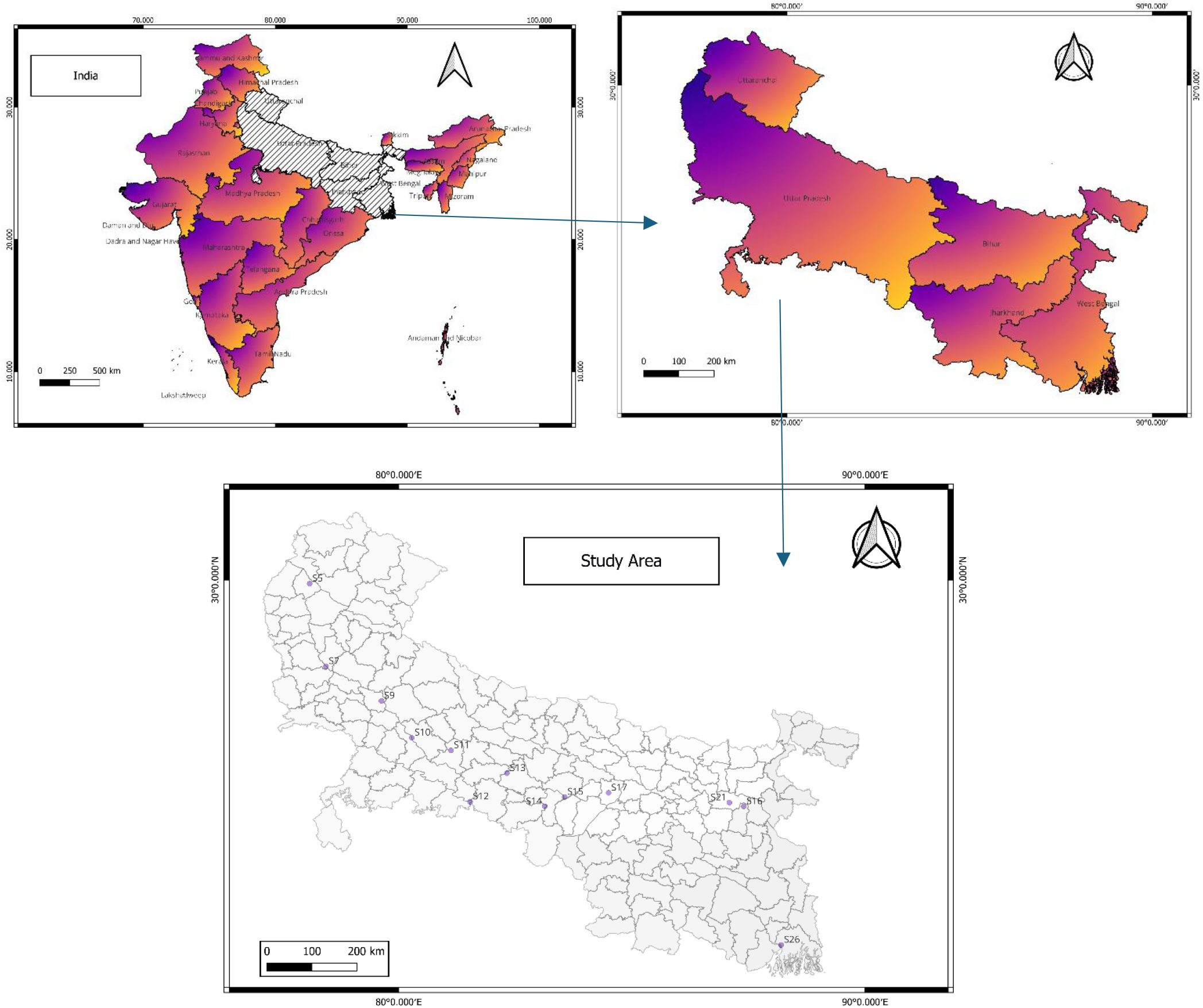
Occurrence of *Ficus racemose* along the study area.

### Collection of Rhizosphere Soil samples

Soil samples from the rhizosphere of *Ficus racemosa* plant were procured from diverse locations spanning the vast geographical expanse from Gaumukh to Gangasagar. Notably, the state of Uttarakhand presented occurrences in Haridwar (S5), while Uttar Pradesh featured locations such as Bijnor (S6), Narora (S7), Badaun (S8), Farrukhabad (S9), Bithoor (S10), Dalmau (S11), Prayaraj (S12), Mirzapur (S13), Varanasi (S14), and Ballia (S15). Additionally, Bihar hosted sites in Revelganj (S16), Patna (S17), Barh (S18), Bahachoki (S19), and Farka (S20). In Jharkhand, the collection site was limited to Sahibganj (S21), while West Bengal encompassed Farraka (S22), Hazarduari (S23), Mayapur (S24), Hooghly (S25), and Gangasagar (S26). The soil sampling procedure involved careful excavation around plant roots to a depth of 15 cm using a tool. Subsequently, a 100-gm wet soil specimen was delicately collected on paper from this depth and placed in an open area for drying under a fan. After 30 minutes, the soil achieved moisture-free status, underwent meticulous debris removal, and fine soil separation from the residue using filter paper.

### Rationale of plant selection

The selection of *Ficus racemosa*, commonly known as the cluster fig tree or gular fig, for study can be justified based on both its abundance and medicinal properties. *Ficus racemosa* is a widely distributed tree species found in various regions, including parts of Asia, Africa, and Australia. Its abundance in these regions makes it readily accessible for research and cultivation, providing ample opportunities for studying its medicinal properties. *Ficus racemosa* possesses several medicinal properties that have been recognized and utilized in traditional medicine systems. Various parts of the tree, including its leaves, bark, fruits, and latex, are used in herbal preparations. The tree’s bark and latex are known for their anti-inflammatory and wound-healing properties, making them valuable in treating skin disorders and injuries. Additionally, extracts from the bark and leaves have been reported to exhibit antimicrobial activity against certain pathogens, further enhancing their medicinal value. Moreover, *Ficus racemosa* is also known for its potential pharmacological effects in managing various health conditions such as diabetes, gastrointestinal disorders, and respiratory ailments. Compounds isolated from different parts of the plant have shown promising results in scientific studies, indicating its potential as a source of novel therapeutic agents.

### Standard Method of Isolation of Rhizosphere Bacteria

The study commenced with the collection of rhizosphere soil samples obtained from the root zones of target plants, ensuring spatial diversity through sampling from multiple locations. Following collection, the samples underwent meticulous processing to eliminate debris and large particles, employing techniques such as sieving or centrifugation to achieve a homogenized soil sample suitable for subsequent analysis. Serial dilution of the processed soil sample was then carried out using sterile diluents to reduce microbial population density to manageable levels, facilitating the isolation of individual bacterial colonies. These diluted samples were plated onto both selective and non-selective agar media using spread plate or pour plate methods, with selective media tailored to specific bacterial groups based on metabolic traits or antibiotic resistance, while non-selective media supported the growth of a broad spectrum of bacteria. Incubation of the agar plates followed, under optimized conditions of temperature and moisture conducive to bacterial growth, with incubation periods tailored to the bacterial species under investigation and the media utilized. Post-incubation, morphologically distinct bacterial colonies were selectively isolated for further analysis, their varying size, shape, color, and texture offering initial insights into the diversity of rhizosphere bacteria. Enumeration of cultivable bacterial populations in the rhizosphere was achieved through the quantification of colony-forming units (CFUs) on agar plates, providing valuable information regarding the abundance and distribution of rhizosphere bacteria within the soil matrix.

### Media used for the bacteria cultivation

Various types of growth media are used for bacterial cultivation, each serving different purposes based on the specific requirements of the bacteria being studied. Nutrient agar serves as a general-purpose medium, providing a rich source of nutrients necessary for the growth of various bacteria. Eosin Methylene Blue Agar (EMB) and MacConkey agar is a selective and differential medium for the isolation of Gram-negative enteric bacteria. Similarly, some selective specific media is used for the cultivation of beneficial bacteria such as Aleksandrow medium is specialized for Mycobacterium tuberculosis, containing specific nutrients and inhibitors tailored for its growth. Zinc Solubilizing Medium isolates bacteria capable of solubilizing zinc from insoluble compounds. Rhizobium medium supports the growth of nitrogen-fixing Rhizobium bacteria, while Azotobacter agar is for isolating and cultivating Azotobacter species, both important for plant health. Azospirillum Medium, without agar, is for liquid culture of Azospirillum species, facilitating experimental procedures. Pikovskaya’s broth cultivates phosphate-solubilizing bacteria, useful in studying their role in promoting plant growth by making phosphate available for uptake. Each medium serves a specific purpose, tailored to isolate and study targeted bacterial species in diverse environments.

### Count of the cultivable organism

The quantification of cultivable bacterial concentration through Colony Forming Units (CFU) involves culture-based counting methods. Serial dilution and spread plating techniques are employed to determine the number of culturable bacteria present per 1 mL of culture (CFU/mL). While this method provides a direct measurement of bacterial cell counts, it tends to undercount the actual number of bacteria, as it only detects those that can grow on specific solid media, excluding unculturable, live, inactive, or damaged bacterial cells. The bacterial concentration was measured using the standard method of the American Society of Microbiology. This involved serial dilutions of 1:10, spreading onto agar plates of the correct growth medium, labeling plates with the Dilution Factor (DF), and incubating overnight. Colony counts were then used to calculate Total CFU/mL, with all calculated CFU values converted to Log10 values for analysis.

### Isolation of active phytoconstituent of the plant

Isolating active phytoconstituents from plants involves several steps aimed at extracting and purifying the bioactive compounds for further analysis or application. The steps are given below:

### Plant material collection

Bulk plant samples for phytochemical analysis are collected by beginning with the clear identification of the target plant species. Representative sampling sites are chosen, and a systematic approach is employed to ensure randomness, with clean and sterilized tools utilized to avoid contamination, and separate tools designated for different plant species. Samples are collected, considering factors such as flowering or fruiting periods. Each sample is labeled with key information, and detailed records of the collection process are maintained. Precautions are taken to avoid contaminants, gloves are worn, and samples are transported in breathable containers. An adequate sample size is determined for statistically meaningful results, and ethical guidelines and permitting procedures are followed if collecting from protected areas. Post-collection, samples are cleaned to remove debris, processing steps are documented, and samples are stored in a cool, dark, and dry environment to preserve phytochemical content. Specific methodologies are referred to, and consultation with experts in plant biology or phytochemistry is sought for field-specific insights.

### Preparation of the extracts

For the preparation of extracts, several methods were employed to determine the content of various compounds. For the assessment of tannin content, a 1-10g sample was mixed with 50 ml Milli-Q water, shaken, and sonicated for 30 minutes, followed by filtration and dilution with 750 ml Milli-Q water. Then, 25 ml of indigo sulphonic acid was added and titrated with 0.1N potassium permanganate solution until a golden yellow color endpoint was reached, with a blank titration performed without the sample. Saponin content was determined by gravimetry, where a 5g sample was mixed with 50 ml of 1:1 methanol:water solvent, refluxed for 1 hour, filtered, and concentrated to dryness after three repetitions. The residue was treated with petroleum ether, followed by methanol and acetone, filtered, dried, and weighed. Total polyphenol content was assessed by incubating a 1 ml sample with 1 ml Folin–Ciocâlteu reagent for 5 minutes, then adding 1 ml of 10% sodium carbonate solution and incubating in the dark for 1 hour. Absorbance was measured at 760 nm using a UV-Visible spectrometer, repeating the process for gallic acid as a reference standard to establish linearity. Similarly, total flavonoid content was determined by incubating a 1 ml sample with 0.4 ml of 10% aluminium chloride, 0.4 ml sodium acetate, and 3 ml ethanol at room temperature for 30 minutes, followed by absorbance measurement at 450 nm using a UV-Visible spectrometer, with quercetin dihydrate used as the reference standard to plot linearity. Quantitative analysis of standard compounds was performed accordingly.

### Quantitative analysis of the standard compounds

The quantitative analysis of standard compounds was conducted using established protocols. High Performance Thin Layer Chromatography (HPTLC) analysis was carried out using a CAMAG HPTLC system equipped with an Automatic TLC Sampler (ATS 4), TLC scanner 4, and TLC visualize, with data processing facilitated by win-CATS software (version 1.4.10). For sample preparation, approximately 1 g of each batch sample was dissolved individually in 10 mL of methanol, subjected to 20 minutes of shaking and sonication, and then centrifuged for 5 minutes at 5000 rpm to obtain clear solutions. Samples were spotted as per their serial numbers mentioned in Table 1. Chromatographic separation was achieved using TLC Silica gel 60 F254 aluminum sheets with a mobile phase composed of ethyl acetate: formic acid: acetic acid: water (10: 1.1: 1.1: 2.3). The saturation time was set to 10 minutes, with a migration distance of 70 mm and a band length of 8 mm. Injection volumes of 25 µL were applied, and visualization was performed at 254 nm and 366 nm prior to derivatization. These meticulous conditions ensured efficient separation and accurate analysis of components present in the samples.

The identification of compounds by Ultra Performance Liquid Chromatography Quadrupole Time-of-Flight Mass Spectrometry (UPLC/MS-QToF) involved the analysis of a test sample of Ficus racemosa (PRF/CHI/1023/1004): UP1.P1.S2. Sample preparation consisted of dissolving 250 mg of the Ficus racemosa raw sample in 5 mL of methanol: water (50:50) followed by 30 minutes of sonication. Subsequently, the solution underwent centrifugation at 10000 rpm for 5 minutes and filtration through a 0.22 µm nylon filter. The UPLC/MS-QToF analysis was conducted using a Xevo G2-XS QToF system (Waters Corporation, USA) equipped with Acquity UPLC I Class and Unifi software. Chromatographic separation utilized a ChromCore 120 C18 column (100 x 2.1 mm, 1.8 µm) with a flow rate of 0.3 mL/min, employing a gradient elution of 0.1 % formic acid in water (mobile phase A) and 0.1 % formic acid in acetonitrile (mobile phase B). The column and sample temperatures were maintained at 30°C and 20°C, respectively, throughout the analysis. Detection was performed using the Xevo G2-XS QToF, with 2 µL of the test solution injected into the UPLC/MS-QToF system, and chromatograms were recorded in both positive and negative ionization modes.

## Results and Discussion

### Cultivation of rhizosphere bacterial population

The cultivation of rhizosphere bacterial populations associated with *Ficus racemose* found from different location. Rhizosphere soil samples were collected from Haridwar (Uttarakhand) to Gangasagar (West Bengal). The collected soil samples were processed to remove debris and homogenized to ensure even distribution of microorganisms. Serial dilutions of the soil samples were plated onto nutrient agar and selective media suitable for promoting the growth of rhizosphere bacteria. Following an incubation period under optimal conditions, bacterial colonies were observed on the agar plates. Individual colonies were selected and streaked onto fresh agar plates to obtain pure cultures of rhizosphere bacteria. Morphological characterization techniques were employed to identify the isolated bacterial strains. The results revealed a diverse population of rhizosphere bacteria associated with *Ficus racemose* as shown in

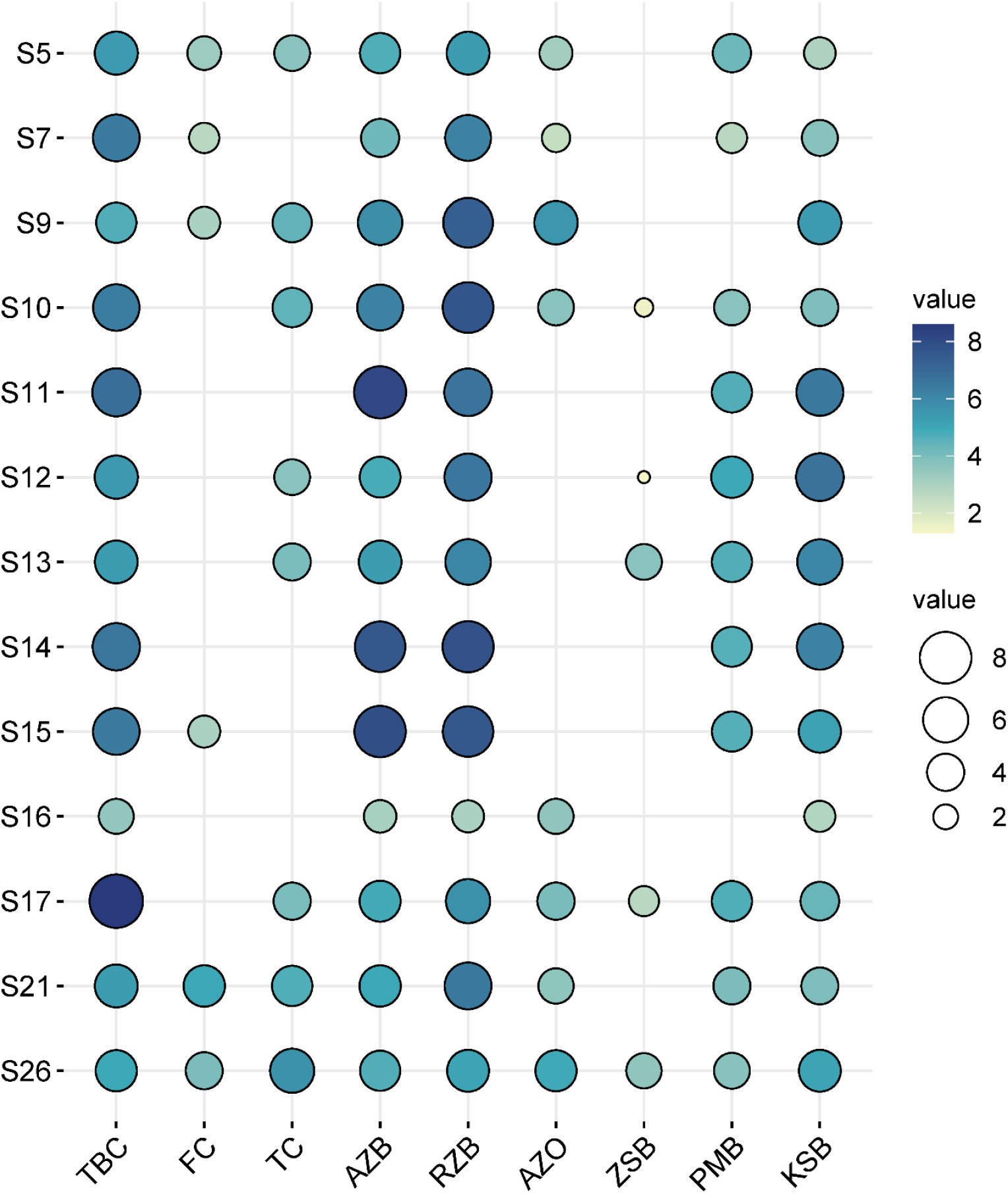

The table presents data on various parameters measured at different sampling sites (S5, S7, S9, S10, S11, S12, S13, S14, S15, S16, S17, S21, and S26), including latitude, longitude, altitude, and microbial counts for total bacterial count (TBC), fecal coliforms (FC), total coliforms (TC), Azotobacter (AZB), Rhizobium (RZB), Azospirillum (AZO), zinc-solubilizing bacteria (ZSB), Potash mobilizer bacteria (PMB), and Phosphate-Solubilizing Bacteria (KSB). Altitude varied among sampling sites, ranging from 3.00 to 265.00 meters above sea level. Geographical coordinates (latitude and longitude) indicated the diverse locations sampled, reflecting variations in environmental conditions. The total bacterial count (TBC) varied across the sites, ranging from 3.54 to 8.60 log CFU/g. Site S17 exhibited the highest TBC, while site S16 had the lowest. Fecal coliforms (FC) and total coliforms (TC) were also measured, indicating potential fecal contamination and overall microbial load, respectively. Site S17 showed the highest FC count, suggesting a possible contamination source in the vicinity. *Azotobacter* (AZB), Rhizobium (RZB), and *Azospirillum* (AZO) counts varied significantly among the sampling sites. Sites with higher counts of these beneficial bacteria, such as S17 and S10, may indicate favorable soil conditions for plant growth and nitrogen fixation. Zinc-solubilizing bacteria (ZSB) counts ranged from 1.30 to 7.83 log CFU/g, indicating the presence of bacteria capable of solubilizing zinc, which is essential for plant nutrition. Pikovskaya’s broth (PMB) counts, used to assess phosphate-solubilizing bacteria, varied across sites, indicating differences in phosphate solubilization potential. The results highlight the heterogeneity of microbial populations across the sampled locations, reflecting differences in soil characteristics, land use, and agricultural practices. High counts of beneficial bacteria such as *Azotobacter, Rhizobium*, and *Azospirillum* at certain sites suggest the potential for enhancing soil fertility and plant growth through microbial inoculation or soil management practices. The presence of fecal coliforms at some sites indicates potential environmental contamination, necessitating remediation efforts to safeguard public health and ecosystem integrity.

### Identification of Phytoconstituent

In the identification of phytoconstituents, we employed a combination of analytical techniques to characterize the chemical composition of the plant extract. Initially, the crude extract was subjected to preliminary screening tests to detect the presence of major classes of phytoconstituents such as tannin, flavonoids, polyphenol and saponins. Subsequently, we employed advanced analytical methods including chromatography (such as high-performance liquid chromatography, HPLC) coupled with mass spectrometry (LC-MS/MS) or UV-Vis spectroscopy to further analyze and identify specific phytoconstituents present in the extract. Chromatographic separation facilitated the isolation of individual compounds, while mass spectrometry provided molecular weight information and fragmentation patterns, aiding in compound identification. Additionally, spectral data obtained from UV-Vis spectroscopy allowed for the determination of characteristic absorption peaks, aiding in the identification of compounds based on their UV profiles. By comparing the obtained data with reference standards, literature databases, and spectral libraries, we were able to identify several phytoconstituents present in the plant extract, including but not limited to flavonoids, phenolic acids, alkaloids, and terpenoids.

According to the data, the tannin content in *Ficus racemosa* is notably elevated in S22 (Farraka), with a value of 0.179. Conversely, S9 (Farrukhabad) and S11 (Dalmau) exhibit the least tannin content, both recording a value of 0.019. The subsequent component under investigation is saponin, with the results revealing that S1 (Gaumukh) exhibits the highest saponin content at 9.43, while the lowest saponin content is observed in S20 (Farka) at 2.33. The analysis of flavonoid content reveals that the highest flavonoid levels are found in S7 (Prayagraj) and S9 (Varanasi), both recording a value of 0.058. Conversely, the lowest flavonoid content is observed in S16 (Revelganj) at 0.005. The assessment of total polyphenol content indicates that S22 (Farraka) has the highest value at 1.446, while the lowest total polyphenol content is recorded in S20 (Farka) at 0.067. Fig.

### Quantitative analysis of the standard compound

The elucidation of standard components via Ultra-Performance Liquid Chromatography/Mass Spectrometry-Quadrupole Time-of-Flight (UPLC/MS-QToF) represents a robust analytical methodology for the resolution, detection, and characterization of compounds within a given sample. In the context of *Ficus racemosa*, a total of 50 compounds were identified using UPLC/MS-QToF. In Positive ionization mode, out of 24 identified compounds the top 5 major components, as determined by concentration, are Quercetin-3-Glucuronide, Racemosic acid, Kaempferol-3-O-glucuronide, Luteolin 7-O-diglucuronide, and Dihydroferulic acid 4-O-glucuronide, as depicted in Fig. 3.6. Conversely, in Negative ionization mode, out of 26 compounds, the the principal components are identified as 7,10,12-trihydroxy-8-octadecenoic acid, Quercetin-3-Glucuronide, Dihydroferulic acid 4-O-glucuronide, 17-Hydroxylinolenic acid, and Racemosic acid, as illustrated in Fig. 3.7. The high-resolution mass spectra and chromatographic separation facilitated by UPLC/MS-QToF contribute to the accurate identification and quantification of these significant components, providing valuable insights into the chemical composition of *Ficus racemosa*.

**Fig. 3.6.**
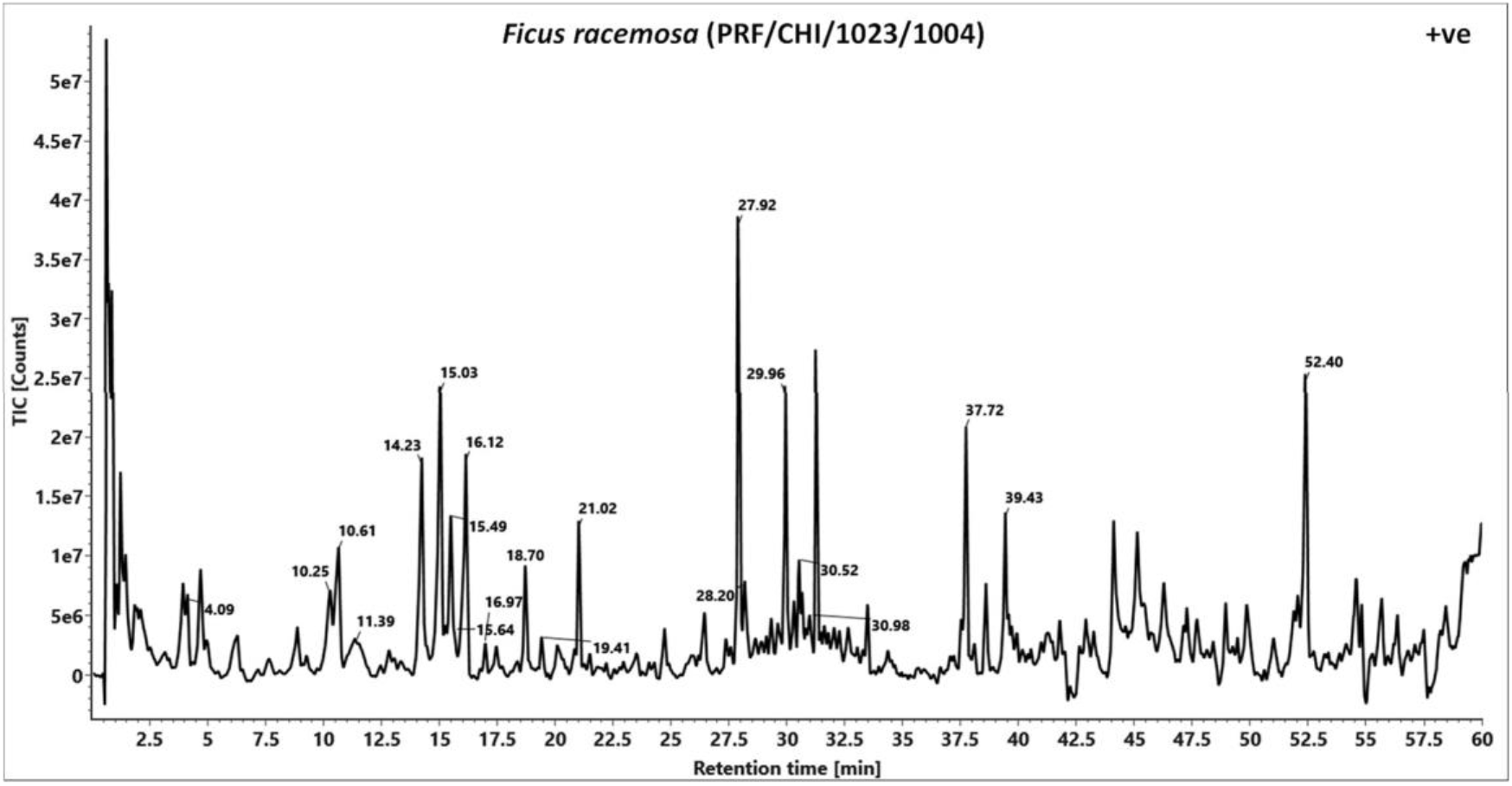
TIC chromatogram of *Ficus racemosa* PRF/CHI/1023/1004) in positive ionizationmode.

**Fig. 3.7.**
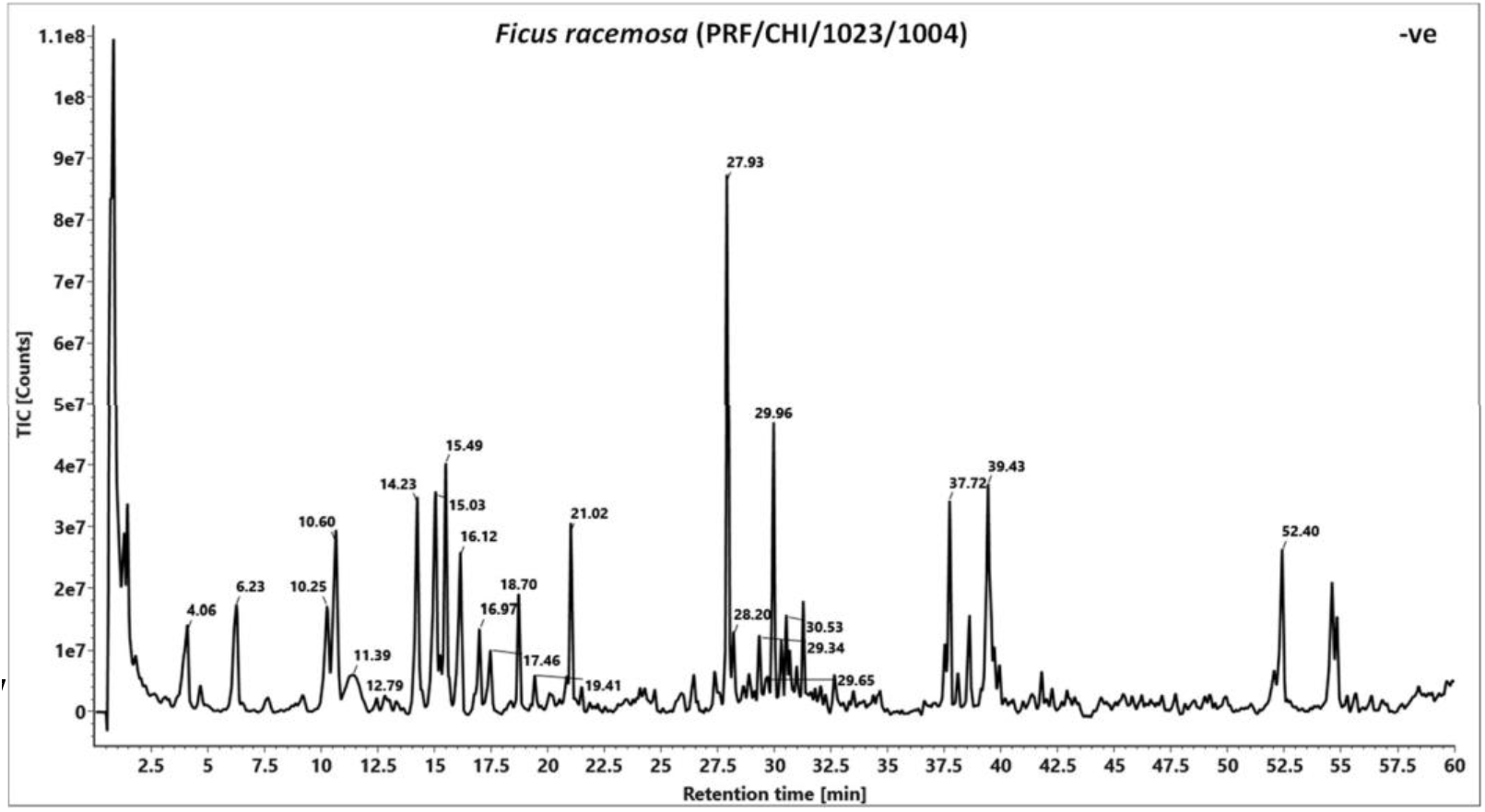
TIC chromatogram of *Ficus racemosa* PRF/CHI/1023/1004) in negative ionizationmode.

### Quantitative analysis of the markers

For the quantitative analysis of the marker aout 1.5 gm of the sample was dissolved in 25 ml of water: methanol (50:50), sonicated for 2 hours at 60 oC and filter through 41 number whatman filter paper. Repeat the extraction one more time, pool all the extract and make up the to 50mL. Filter through 0.45 µ filtr paper, and use for the analysis and Prepared Mix standard of Neo chlorogenic acid, Chlorogenic acid and Crypto Chlorogenic acid of 50 ppm from 1000 ppm stock solutions. Analysis was performed on Prominence-i HPLC system (Shimadzu, Japan). Separation was achieved using a Shodex C18-4E (5 µm, 4.6*250 mm) column subjected to binary gradient elution. The two solvents used for the analysis were water containing 0.1% Acetic acid in water (solvent A) and acetonitrile (solvent B). Column temperature was kept 35oC and flow was set 1.0 mL/min during the analysis. 10 µL of standard and test solution were injected. Wavelength set at 325 nm (for Neochlorogenic acid, Chlorogenic acid and Cryptochlorogenic acid).

In the HPLC analysis, the most substantial concentration of Neochlorogenic acid is observed in S9 (Farrukhabad), quantified at 0.048. In S13 (Mirzapur), Chlorogenic Acid is detected, registering a measured quantity of 0.025. Furthermore, S9 (Farrukhabad) exhibits the highest content of Cryptochlorogenic acid, identified with a recorded amount of 0.025. The mean concentrations of Neochlorogenic acid, Chlorogenic Acid, and Cryptochlorogenic acid across all samples are determined as 0.0174, 0.0259, and 0.012667, respectively.

### Discussion regarding what is the possible explanation of the relationship

The relationship observed between rhizosphere bacterial populations and phytoconstituents in the discussed study can be explained by several underlying mechanisms Rhizosphere bacteria play crucial roles in nutrient cycling and soil health. Beneficial bacteria such as Azotobacter, Rhizobium, and Azospirillum are known for their ability to fix atmospheric nitrogen, solubilize phosphorus, and produce plant growth-promoting substances like phytohormones. These activities enhance nutrient availability to plants, promoting their growth and metabolism, including the synthesis of phytoconstituents. Rhizosphere bacteria can indirectly influence plant physiology by modulating soil properties and microbial communities. For instance, zinc-solubilizing bacteria contribute to zinc availability in the soil, which is a cofactor for many enzymes involved in plant secondary metabolite biosynthesis. Similarly, phosphate-solubilizing bacteria enhance phosphate availability, a key component in energy metabolism and secondary metabolite biosynthesis. Plants and rhizosphere bacteria engage in intricate signaling and communication processes. Beneficial bacteria can induce systemic responses in plants, triggering the activation of defense mechanisms and secondary metabolite production. This phenomenon, known as induced systemic resistance (ISR) or systemic acquired resistance (SAR), can enhance the synthesis of phytoconstituents involved in plant defense against pathogens and herbivores. Rhizosphere bacteria can mitigate abiotic stresses in plants, such as drought, salinity, and heavy metal toxicity. By enhancing plant stress tolerance, these bacteria can alleviate the negative impacts of environmental stressors on plant metabolism, allowing plants to allocate resources towards secondary metabolite production. The composition and diversity of rhizosphere microbial communities can directly influence plant secondary metabolism. Certain bacterial taxa may directly stimulate or inhibit the synthesis of specific phytoconstituents through direct interactions with plant tissues or by modulating microbial community dynamics in the rhizosphere. Overall, the relationship between rhizosphere bacterial populations and phytoconstituents is multifaceted and influenced by various factors including nutrient availability, plant-microbe interactions, and environmental conditions. Understanding these mechanisms is essential for harnessing the potential of rhizosphere bacteria to enhance plant productivity, health, and the production of bioactive compounds with agricultural and pharmaceutical applications.

## Conclusion

In conclusion, our research sheds light on the intricate relationship between rhizosphere bacterial concentration and phytoconstituents in *Ficus racemosa*. Through comprehensive analysis across various sampling sites, we have observed substantial heterogeneity in rhizosphere bacterial populations, indicative of diverse soil conditions and microbial communities. Notably, sites exhibiting higher counts of beneficial bacteria, including Azotobacter, Rhizobium, and Azospirillum, corresponded to elevated levels of phytoconstituents such as flavonoids, tannins, and total polyphenols. This correlation underscores the potential influence of rhizosphere bacteria on plant secondary metabolism, possibly mediated through mechanisms such as nutrient availability, stress alleviation, and symbiotic interactions. Moreover, our identification of specific phytoconstituents and their spatial distribution provides valuable insights into the chemical diversity of *Ficus racemosa* extracts. Variations in phytoconstituent levels among sampling sites highlight the complex interplay between environmental factors, microbial communities, and plant metabolism. HPLC analysis further delineates distinct concentrations of bioactive compounds, underscoring the importance of considering spatial variability in phytochemical profiles.

Overall, our findings contribute to a deeper understanding of plant-microbe interactions in the rhizosphere and their implications for plant health, soil fertility, and ecosystem functioning. By elucidating the role of rhizosphere bacteria in modulating plant secondary metabolism, our research offers avenues for harnessing microbial communities to enhance phytochemical diversity and promote sustainable agriculture. Further investigations into the molecular mechanisms underlying these interactions will pave the way for innovative strategies in agricultural and pharmacological applications.

